# ISTDECO: In Situ Transcriptomics Decoding by Deconvolution

**DOI:** 10.1101/2021.03.01.433040

**Authors:** Axel Andersson, Ferran Diego, Fred A. Hamprecht, Carolina Wählby

## Abstract

In Situ Transcriptomics (IST) is a set of image-based transcriptomics approaches that enables localisation of gene expression directly in tissue samples. IST techniques produce multiplexed image series in which fluorescent spots are either present or absent across imaging rounds and colour channels. A spot’s presence and absence form a type of barcoded pattern that labels a particular type of mRNA. Therefore, the expression of a gene can be determined by localising the fluorescent spots and decode the barcode that they form. Existing IST algorithms usually do this in two separate steps: spot localisation and barcode decoding. Although these algorithms are efficient, they are limited by strictly separating the localisation and decoding steps. This limitation becomes apparent in regions with low signal-to-noise ratio or high spot densities. We argue that an improved gene expression decoding can be obtained by combining these two steps into a single algorithm. This allows for an efficient decoding that is less sensitive to noise and optical crowding.

We present IST Decoding by Deconvolution (ISTDECO), a principled decoding approach combining spectral and spatial deconvolution into a single algorithm. We evaluate ISTDECO on simulated data, as well as on two real IST datasets, and compare with state-of-the-art. ISTDECO achieves state-of-the-art performance despite high spot densities and low signal-to-noise ratios. It is easily implemented and runs efficiently using a GPU.

ISTDECO implementation, datasets and demos are available online at: github.com/axanderssonuu/istdeco

## 1 Introduction

Identification of cell types in a tissue provides invaluable information on biological processes such as development and disease progression. Cells can be identified based on transcriptomics, i.e., their gene expression. Single-cell RNA sequencing techniques (Svensson *et al.*, 2018; Grün and van Oudenaarden, 2015) make it possible to identify cell types and monitor the heterogeneity of complex tissues. To understand the functional architecture of a tissue it is essential to reconstruct the spatial organisation of its constituent cell types. Therefore, single cell sequencing analyses are often complemented with imaging-based methods for in situ spatial transcriptomics, or IST (Ke *et al.*, 2013; Shah *et al.*, 2016; Wang *et al.*, 2018; Moffitt *et al.*, 2016; Codeluppi *et al.*, 2018; Eng *et al.*, 2019). These methods enable mapping of gene expression in the form of mRNA molecules directly in tissue samples. This in turn enables identification of specific cell type location, so that the functional roles of cells inside the tissue architecture can be explored. Typically, IST techniques produce large image series where small but bright spots are present and absent in different colour channels and imaging rounds, see Figure 1a. The spots themselves originate from fluorophores that have bound to different parts of the mRNA sequences. The fluorophores start to fluoresce when irradiated with light of a particular wavelength, making it possible to image the molecules. Due to diffraction and aberration in the microscope, the imaged fluorophores are blurred by a point spread function that is often approximated by a Gaussian. The combination of absent and present spots across rounds and channels, at a particular location, forms an on-off intensity pattern, i.e., a type of barcode. IST experiments are usually designed such that different types of barcoded patterns are observed for different types of mRNA molecules. The location and mRNA type can therefore be determined by localising the Gaussian shaped spots and decode the barcoded intensity pattern across the rounds and channels. This is usually performed automatically using different types of image analysis pipelines.

**Fig. 1.**
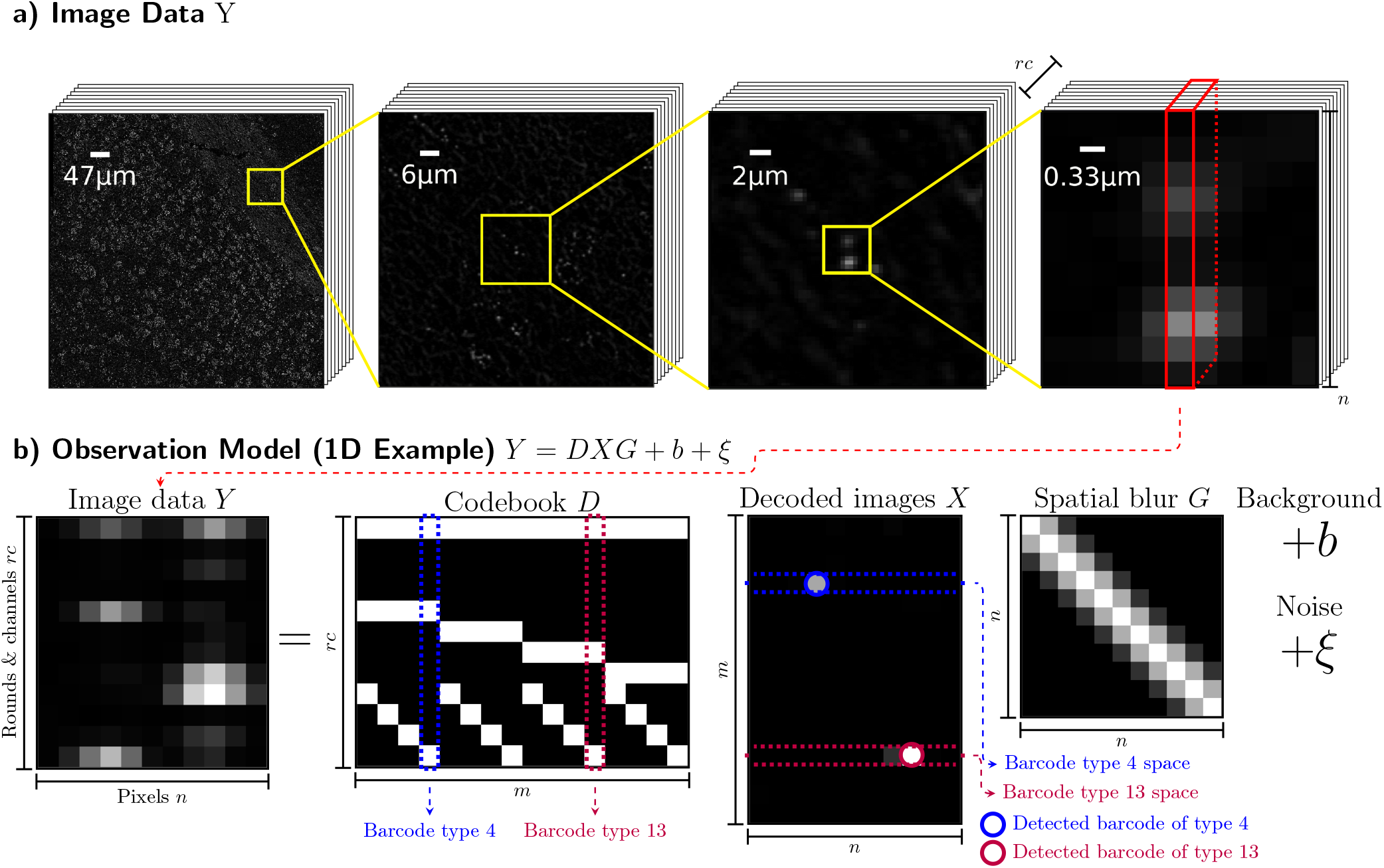
**(A)** Example of image data from an IST experiment with *r* sequencing rounds and *c* colour channels. Fluorescent spots are spread out across the tissue and are present and absent across imaging rounds and colour channels. Zooming in reveals the Gaussian shape of the signal spots. **(B)** Given observations *Y*, the idea of ISTDECO is to find an *X* such that *Y* = *DXG* + *b* + *ξ*, where *D* is a known codebook with targeted barcode types, *G* is a known blurring kernel that models the point-spread-function, and *b* is known background. The variable *X*, referred to as the decoded images, tells which barcode type is present at what location.

### 1.1 Related Work

Ke *et al.* (2013) introduced one of the first image analysis pipelines for IST experiments. Here, spots are first located and segmented in a reference image. This reference image is obtained by using an anchor sequence that simultaneously labels all targeted mRNA. A spot’s location is then defined as the centre of the segment. The mRNA type is decoded by matching the observed barcode with a list of targeted barcode types — each type corresponding to an mRNA type. This list is often referred to as a *codebook*. Another algorithm was presented by Chen *et al.* (2015). Here, signal spots are first localised in individual channels using a Gaussian-fitting algorithm (Babcock *et al.*, 2012). This allows partially overlapping spots to be localised. The localised spots are connected across different rounds based on spatial proximity to create a barcode that is decoded using the codebook. To process higher volumes of data, the pipeline was later modified by Moffitt *et al.* (2016). In this pipeline individual pixels are first labelled with a barcode type from the codebook. Adjacent pixels with the same label are called as the same RNA.

The problem of optical crowding was also addressed by Codeluppi *et al.* (2018) who proposed a non-barcoded approach. Here, individual spots are enhanced using Laplacian of Gaussian (LoG) filters and detected as local maxima above an automatically calculated threshold. The LoG filters were also used by Wang *et al.* (2018) to localise the fluorescent spots. The most dominant colour in each sequencing round is used to create a barcode that is decoded using a codebook. An initiative was made in 2017 to unify the image analysis techniques of the different IST decoding methods into a single framework. The result is an open source Python based library named Starfish (Perkel, 2019; Shannon *et al.*, 2018). Today, Starfish provides decoding pipelines for IST methods such as (Ke *et al.*, 2013; Moffitt *et al.*, 2016; Codeluppi *et al.*, 2018; Wang *et al.*, 2018). The decoding pipelines are inspired by the pipelines from the original publications but are adapted into the Starfish framework.

The most closely related method — developed completely independently from ours — is BarDensr (Chen *et al.*, 2020), which relies on a nonnegative matrix regression for unmixing and deconvolving multiplexed image data. BarDensr also models physical properties such as signal gain, phasing, and spectral overlap, and builds on optimising a sparsity regularised least squares problem.

### 1.2 Our contribution

Summarising existing IST techniques, we notice that methods either first locate spots and then decode the barcode (Ke *et al.*, 2013; Chen *et al.*, 2015; Wang *et al.*, 2018), or first decode the barcode and then determine the location of the spots (Moffitt *et al.*, 2016). We also note that recent IST techniques allow for larger gene panels as well as increased measurement throughput (Moffitt *et al.*, 2016), but are limited by optical crowding (Chen *et al.*, 2015; Codeluppi *et al.*, 2018). Although existing algorithms are efficient, they are fundamentally limited by strictly separating the localisation and decoding steps. This limitation becomes especially apparent in regions with low signal-to-noise ratio or high spot densities. We argue that an improved spot localisation and barcode decoding can be obtained by **simultaneously** solving the two problems. We summarise our contribution as follows:

- We propose a tool named In Situ Transcriptomics Decoding by Deconvolution (ISTDECO) which simultaneously deconvolves the barcode type and location from multiplexed IST images. The result is a set of non-multiplexed images where individual sparse spots directly indicate the location of a particular type of mRNA. This dramatically simplifies the gene expression quantification.
- We show that deconvolution-based approaches maintain high precision and recall despite low signal-to-noise ratios and high spot densities, achieving state-of-the-art performance.
- By evaluating ISTDECO on data from two real IST experiments, we conclude that barcodes detected with ISTDECO are highly correlated with barcodes detected with two state-of-the-art algorithms.
- We make the tool freely available at github.com/axanderssonuu/istdeco.

## 2 Methods

### 2.1 Notation & Intuition

To describe ISTDECO we start by introducing a few matrices. We let our image data be denoted by 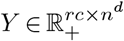, where *r* is the number of staining rounds, *c* is the number of colour channels, *n* is the image width, and *d* is the number of spatial dimensions. Next, we model the spatial shape of the signal spots using a Gaussian kernel parameterised by the standard deviation *σ*_psf_. To conveniently be able to use vectorised notation, we represent convolution with this kernel in terms of a matrix 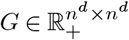. The spots are also present and absent across different rounds and channels, hence forming different types of barcoded patterns. The different types of barcoded patterns are collected in a codebook 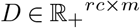, where *m* is the number of different barcode types. With these matrices we stipulate the observation model

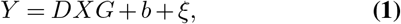

where 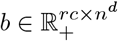 corresponds to background and *ξ* represents noise. The matrix 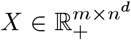 contains coefficients that indicate which barcode type are present at each spatial location. That is, *X* can be interpreted as a reshaped image with *m* different channels, each channel corresponding to a particular barcode type. The channels show small point-like spots that indicate the location of the different barcode types. Determining the matrix *X* therefore dramatically simplifies the localisation and decoding of the gene expression as an individual spot in *X* would directly indicate the location and barcode type. This is exemplified in Figure 1b. As *Y*, *D* and *G* are known, and *b* can be estimated using standard image processing, a naive approach is to directly solve *X* from Eq. 1. Unfortunately, the equation is ill-conditioned. However, we can exploit the prior knowledge that *X* should be nonnegative. Hence an alternative approach is to instead estimate a nonnegative *X* using constrained optimisation. In practice this means finding the minimiser of a function that measures the difference between *Y* and *DXG* subjected to constraints on nonnegativity. Next, we derive this function using maximum likelihood estimation (MLE).

### 2.2 The Objective Function

To derive our objective function, we start by assuming a setting with Poisson distributed noise. We claim that this is an adequate assumption as the pixels in the camera effectively count the number of photons emitted from the fluorophores. The photon count usually follows a Poisson statistic. Mathematically, we assume Poisson distributed noise with mean *DXG* + *b*. The MLE of such a distribution is obtained by maximizing

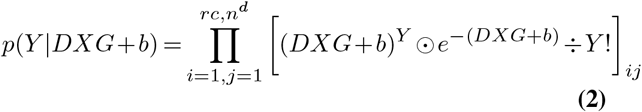

with respect to *X*. The symbol denotes ꖴ the Hadamard product and the divisions, exponents, factorials, and exponentials are elementwise. In practice, maximising Eq. 2 is numerically inconvenient, and it is better to minimise the negative logarithm of the likelihood. Dropping all terms independent of *X*, we have to solve

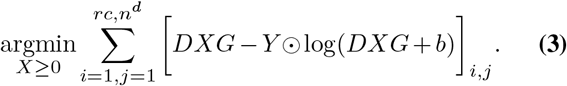

### 2.3 Minimising the objective function

To find a nonnegative *X* we follow Lee and Seung (2001) and derive the following update rule:

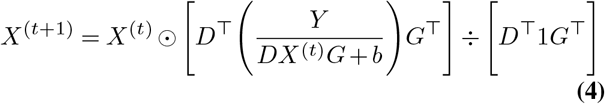

where 1 is an *rc* × *n^d^* matrix with ones. This update is commonly used in nonnegative matrix factorisation algorithms (Lee and Seung, 2001) as well as in Richardson-Lucy deconvolution (Bertero *et al.*, 2009). It possesses a few implicit biases. Firstly, consecutive updates converge to the MLE. Secondly, if all matrices are initialised as nonnegative then *X* will remain nonnegative over the updates. Thirdly, if *X*^(*t*+1)^ is obtained from Eq. 4, and if *b* = 0, then

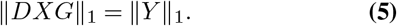

Such a constraint is known to induce sparsity (Bertero *et al.*, 2009) and may seem intuitive in the sense that “no fluorescent signal is thrown away”.

### 2.4 Simultaneous localisation and barcode decoding

Deconvolving the image data through iterations of Eq. 4 enables the simultaneous localisation and decoding. This is done by performing a spatial non-maxima suppression in each of the channels in *X* and picking signal spots with intensities above a threshold *τ*_s_. Furthermore, we occasionally observed barcoded intensity patterns that were not present in the codebook (possible a result of uneven washing between sequencing rounds or cross-reactivity of probes). In those situations, a combination of coefficients in *X* were used to explain the observed pattern, resulting in false-positive detections. To filter these detections, we introduced a barcode quality feature which compares explained with observed signal:

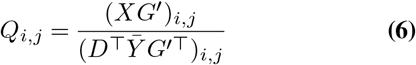

where 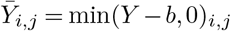 and the multiplication with *G′* corresponds to blurring with a constant filter with width equal to 2[3*σ*_psf_]+ 1. The set of decoded barcodes is thus

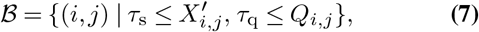

where *i* indicates the barcode type, *j* the barcode location, *X′* is the non-max suppressed *X*, and *τ*_q_ is a threshold for the quality feature. The parameters *τ*_s_ and *τ*_q_ must be tuned empirically. For this matter we found it practical to include a set of nontargeted barcodes in the codebook. The number of nontargeted barcodes detected can help us assess the false discovery rate (FDR), and help us tune the thresholds. We define this FDR as:

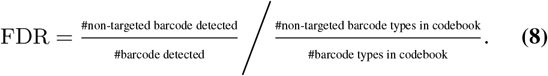

The FDR is 1 when the method randomly assigns barcode types from the codebook and 0 when no nontargeted barcodes are detected. In practise, we found it convenient to initially set thresholds to be very inclusive, and during post-processing tune the thresholds such that a desired FDR is obtained.

## 3 Experiments on Synthetic Data

We first evaluate the proposed decoding method on synthetic data.

### 3.1 Generating synthetic data

We consider an experiment with *c* = 4 channels and *r* = 5 sequencing rounds, and a codebook of *m* = 100 barcode types. We sample spot locations from a continuous uniform distribution but avoid locations along image edges. Spot intensities are also sampled from the uniform distribution 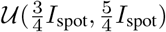. In our experiments we test the decoding on different values for *I*_spot_ and on images with different numbers of barcodes. We also add noise to our image: Noise sampled from a Poisson distribution with *λ*_noise_ = 2 to emulate the stochastic nature of photons interacting with the pixel sensors, and noise from a normal distribution 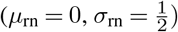 to emulate the noise induced by converting electrons into a digital signal. The shape of the spots are approximated by Gaussians with *σ*_spot_ = 1.2. Figure 2 shows four synthetic images with different numbers of RNA and different signal intensities.

**Fig. 2.**
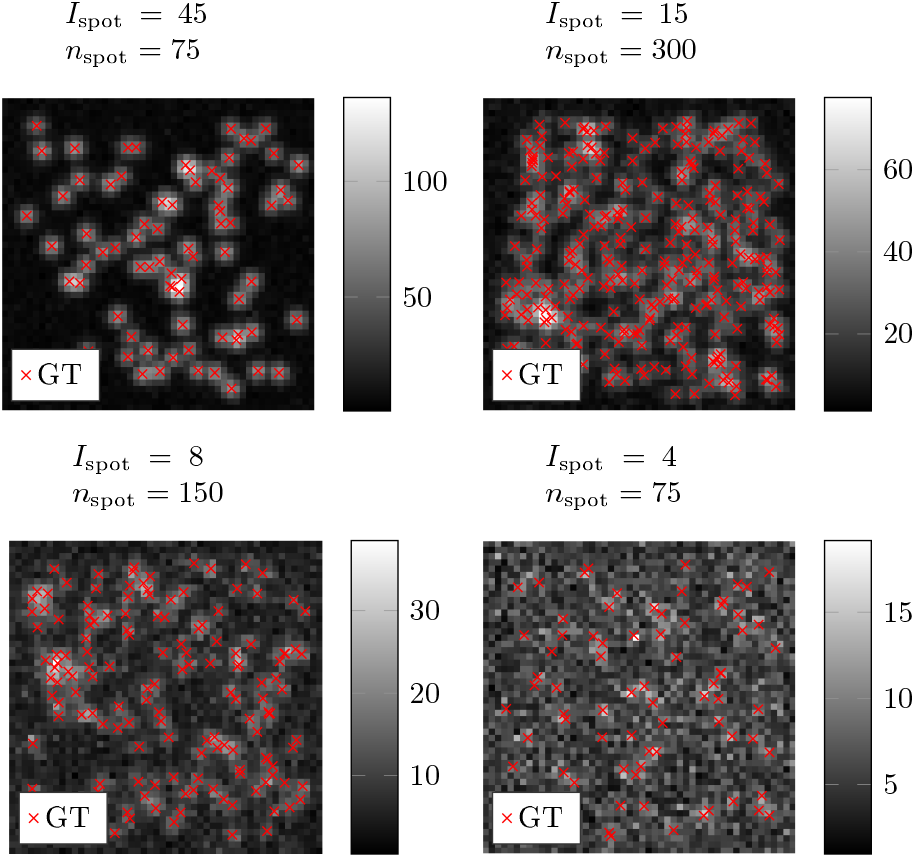
Synthetic images with markers for ground truth (GT) at different signal densities and signal intensities. The images are maximum projected over one sequencing round.

### 3.2 Starfish Decoders

We compare our proposed method with four decoders available in the Starfish library as well as with BarDensr (Chen *et al.*, 2020).

#### *BarDensr* (BarDensr)

BarDensr solves the barcode matching and localisation problem jointly by optimising a sparsity regularised least squares objective function.

#### PixelSpotDecoder (PSD)

A Starfish adaptation of the decoding pipeline presented by Moffitt *et al.* (2016). Individual pixels are decoded into targeted barcodes by comparing the intensity distribution across rounds and channels with a pre-defined codebook. Connected pixels that are mapped to the same barcode are called as the same mRNA.

#### TrackPyDecoder (TPD)

Spots are detected using the Crocker-Grier algorithm (Allan *et al.*, 2014; Crocker and Grier, 1996) from a reference image showing all signal spots. The barcode is determined by taking the maximum across channels in each round at the signal location.

#### LocalMaxPeakFinder (LMPF)

Spots are located as local maxima whose absolute intensities exceed an automatically calculated threshold. The barcode is determined similarly to TPD.

#### BlobDetector (BD)

Spots are detected using multiple Laplacian of Gaussians. Detected spots are connected across rounds and channels, forming a trace which is mapped to a particular barcode. The barcode is determined similarly to TPD.

We also introduced a background removal hyperparameter. For the Starfish decoders, the synthetic images were pre-processed by subtracting this parameter and clipping negative values to zero. For ISTDECO and BarDensr, this parameter was used in the observation model, i.e., as the background offset *b*. Hyperparameters for the respective decoders were tuned using hyperparameter optimisation package HyperOpt (Bergstra *et al.*, 2013).

### 3.3 Evaluation

We evaluate the performance of our proposed method on images with different signal-to-noise ratios (SNR) and on images with different numbers of spots. The different hyperparameters for each method were tuned tuned on a set of five synthetic images and then tested on 30 images. Each method was tuned and tested on the same set of images. The tuning and testing process was repeated for images with different *I*_spot_ as well as images of different numbers of mRNAs. We use the F-measure, (*F*_1_), to evaluate the decoding. A true-positive (TP) is defined as a correctly decoded barcode with distance less than 3 [px] away from a ground truth barcode. A false-positive (FP) is either a wrongly decoded barcode, or a barcode that is more than 3 [px] away from a ground truth barcode. Furthermore, a false-negative (FN) is defined as a ground truth barcode that does not have a correct decoded barcode within 3 [px]. Detected barcodes were paired with ground truth barcodes using the Hungarian algorithm with an assignment reward for true-positive detections.

### 3.4 Results

Figure 3 shows the *F*_1_ measure for the respective methods computed on images with different SNR and images with different numbers of spots. Precision and recall of the different approaches are available in the supplementary information.

**Fig. 3.**
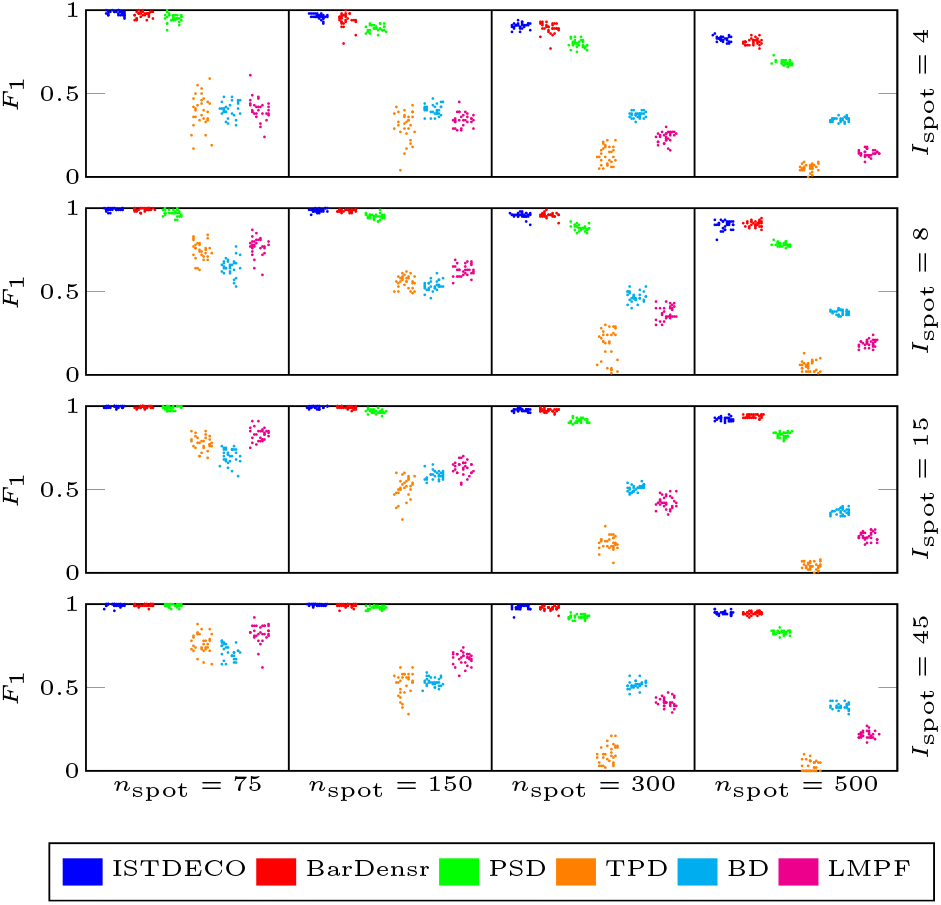
*F*_1_ score computed on image series with different number of spots, *n*_spot_, and different spot intensities, *I*_spot_. The dots have been jittered horizontally to avoid overplotting.

As ISTDECO is implemented solely using vectorised operations it can run very efficiently on a GPU. In Figure 4 we compare the *F*_1_ score of ISTDECO and BarDensr versus wall-clock time. The experiment was carried out on simulated 2D images with *r* = 5, *c* = 4, *n* = 256, *m* = 75, σ_spot_ ~ 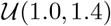. We consider two different signal-to-noise ratios; high SNR (*I*_spot_ = 45), and low SNR (*I*_spot_ = 4), as well as two different spot densities; sparse (*n*_spot_ = 10^3^) and dense (*n*_spot_ = 10^4^). Hyperparameters were tuned using grid search. Details on parameter tuning are available in the supplementary information.

**Fig. 4.**
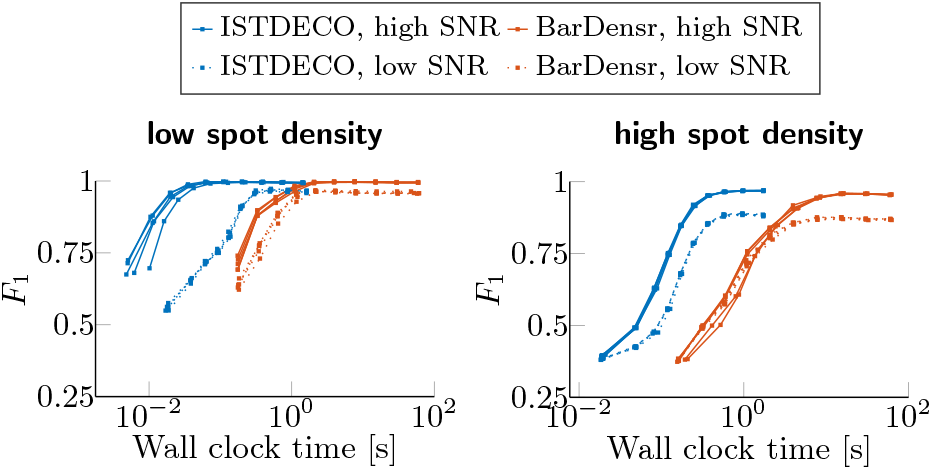
Both ISTDECO and BarDensr decode the gene expression by iteratively optimising an objective function. Consequentially, there is a relationship between the quality of the decoded barcodes and the duration of the optimisation. Here we show the *F*_1_ score versus wall-clock time for the two respective methods on images with different signal-to-noise ratios and spot densities. Both are executed on a GPU.

## 4 Experiments on Real Data

We also consider data from two real IST experiments, and compare ISTDECO with the authors’ original decoding pipelines.

### 4.1 MERFISH Dataset

We first consider a MERFISH dataset, publicly available in Starfish (Shannon *et al.*, 2018), with barcodes localised and decoded using the authors’ original pipeline (Moffitt *et al.*, 2016). We will refer to these detected barcodes as the MERFISH reference barcodes. The image data consists of a 2048 × 2048 [px] field of view with 8 imaging rounds and 2 channels. The codebook comprises 130 different barcode types corresponding to different mRNA species. The codebook also contains 10 different non-targeted barcode types that are used for false-discovery diagnostics. To make the comparison as fair as possible, we utilise the same pre-processing and post-processing strategy as *Moffitt et al.* (2016): First, we pre-process the images using a Gaussian high-pass filter with standard deviation set to 2 [px], and normalise the colour channels using provided scale factors. Thereafter, we follow Moffitt *et al.* (2016) and filter the reference barcodes whose contiguous area was less than 4 [px]. Each of the reference barcodes also have a total magnitude attribute. This attribute reflects the brightness of an observed barcode. We use this attribute in a final post-processing step to filter barcodes such that a pre-selected FDR is obtained.

### 4.2 In Situ Sequencing Dataset

The second dataset that we consider is from an In Situ Sequencing (ISS) experiment (Qian *et al.*, 2020). The dataset is publicly available from Andersson *et al.* (2021) and consists of a 9216 × 4098 [px] large field of view, with 5 rounds and 4 channels, and a codebook consisting of 170 barcodes. Here we have added 9 additional non-targeted barcodes to the authors’ original codebook to allow for false discovery diagnostics. The images are pre-processed with a top-hat filter and registered similar to Qian *et al.* (2020). Finally, we generated a set of ISS reference barcodes using the pipeline by Qian *et al.* (2020). The barcodes were post-processed similarly to Qian *et al.* (2020), i.e., by discarding detected barcodes whose cosine angle to the best matching barcode type in the codebook falls below a threshold. We set this threshold such that a pre-selected FDR is obtained.

### 4.3 Running ISTDECO

We run ISTDECO on the MERFISH image data, as well as the ISS image data for 50 iterations to generate *X*. A Gaussian-shaped point-spread-function with *σ*_psf_ = 1.0 and *σ*_psf_ = 1.75 was used on the respective MERFISH and ISS datasets. We used the provided codebooks to define the codebook matrices. On the ISS images, the cross-talk compensated codebook was used to define *D*. The barcodes in *X* were located after non-max suppressing with a radius of 1.5 [px]. Barcodes with an intensity less than the 99^th^ percentile of the image data were discarded.

### 4.4 Results

After running ISTDECO on the MERFISH and ISS dataset, we compared detected barcodes with those in the reference datasets. Here there are three questions that we want to investigate. Firstly, how many nontargeted barcodes do we detect? Secondly, are the barcodes detected by ISTDECO of the same type as the reference barcodes? Thirdly, are the barcodes detected by ISTDECO located in proximity to the barcodes in the reference data?

We started by investigating the first question. By filtering the barcodes based on the quality score we can obtain different FDR. Figure 5a shows the number of targeted barcodes and number of nontargeted barcodes detected versus different thresholds on the barcode quality score. Furthermore, Figure 5b shows the number of targeted barcodes detected versus FDR rate. Similarly, Figure 5c shows the number of targeted barcodes versus number of nontargeted barcodes detected. If the detected barcodes are filtered such that a very low FDR is obtained, ISTDECO finds roughly the same number of targeted barcodes as the reference methods. However, if we allow slightly higher FDR, then ISTDECO tends to find more targeted barcodes than the reference.

**Fig. 5.**
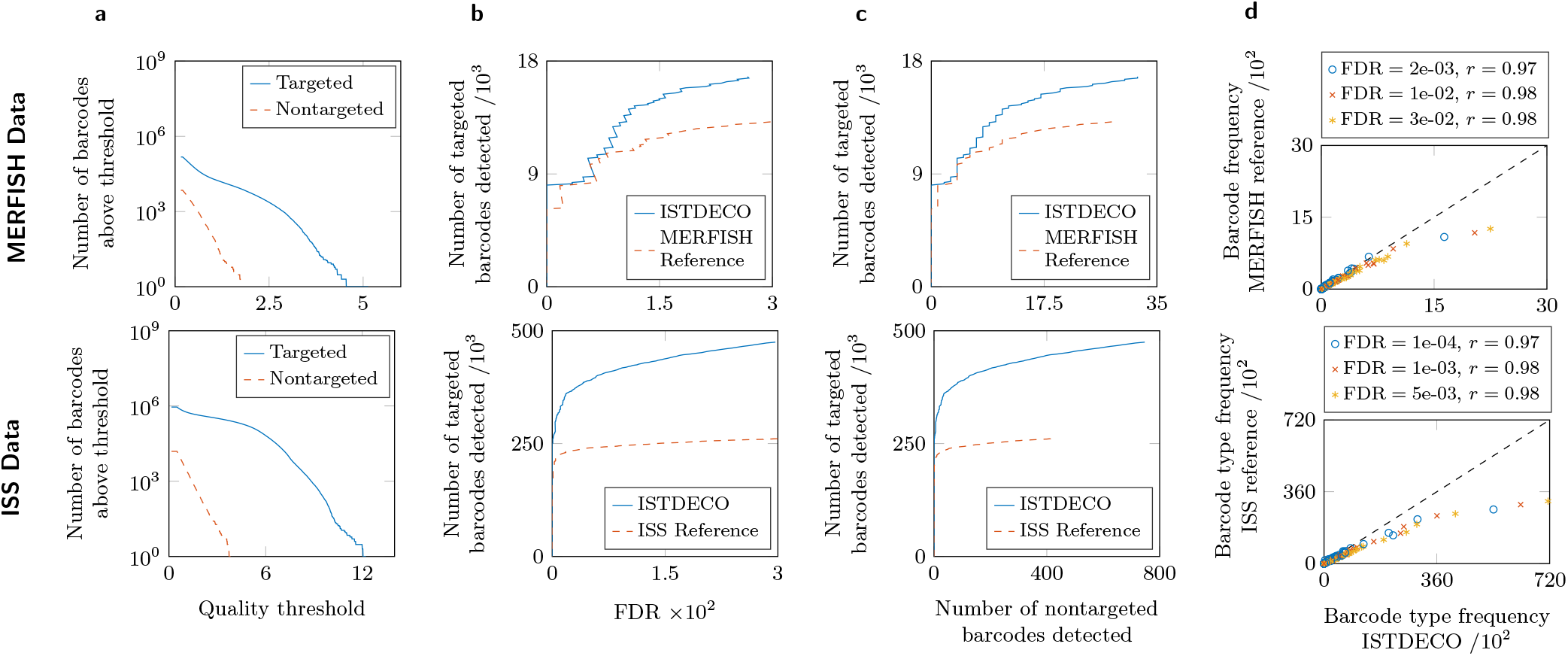
**(A)** The number of targeted and non-targeted barcodes detected by ISTDECO is shown versus the barcode quality threshold. **(B)** Number of targeted barcodes detected is shown versus FDR. The curves are obtained by removing barcodes below the different thresholds, as described in the text. **(C)** The number of targeted barcodes is shown versus the number of nontargeted barcodes. **(D)** The number of barcodes of a particular type detected by ISTDECO is shown versus the number of barcodes of the same type in the reference. This is repeated for three different FDR. *r* corresponds to the Pearson correlation coefficient.

Next, we investigated whether ISTDECO’s detected barcodes are of the same type as the barcodes in the reference datasets. We started by considering three different FDR. For the MERFISH dataset we chose the FDRs 2e-3, 1e-2, 3e-2. Likewise, for the ISS dataset, we chose the FDRs 1e-4, 1e-3 and 5e-3. For a particular FDR, we compared the number of barcodes of each specific type detected by ISTDECO with the number of barcodes of the same types detected by the reference methods, see Figure 5d. Here we see a correlation in the number of barcodes detected, indicating that ISTDECO detects barcodes of a similar type as the reference methods. We use the Pearson correlation coefficient, *r*, to quantify this correlation. For both datasets *r* was approximately around 0.98 for the three considered FDR.

Finally, we wanted to see if the barcodes detected by ISTDECO are spatially close to the reference barcodes. For the MERFISH experiments, we considered two FDRs: A strict rate of 2e-3 and a more inclusive rate of 3e-2. We then computed the spatial distances from each barcode detected by ISTDECO, filtered to obtain the strict FDR, to the nearest barcode of the same type among the MERFISH barcodes, filtered to obtain the inclusive FDR. We found that roughly 80 % of the barcodes detected by ISTDECO were within 3 [px] to a MERFISH reference barcode. We did the opposite as well, i.e., computed the nearest neighbour distance from MERFISH barcodes to ISTDECO barcodes. Figure 6 shows the percentage of detected barcodes that have a nearest neighbour distance less than *d* [px] to a similar barcode, detected using a different method. Here we see that most of the barcodes in the MERFISH and ISS reference data are spatially close to a barcode detected by ISTDECO. However, as the nearest neighbour distance is lower from ISTDECO to one of the reference methods, than from a reference method to ISTDECO, we hypothesize that ISTDECO detects barcodes that are not found by the reference method. Visual assessment of these barcodes indicate that they may be true signals that are missed by the reference method. We refer to supplementary information for figures of barcodes that are only detected by ISTDECO, as well as barcodes not detected by ISTDECO.

**Fig. 6.**
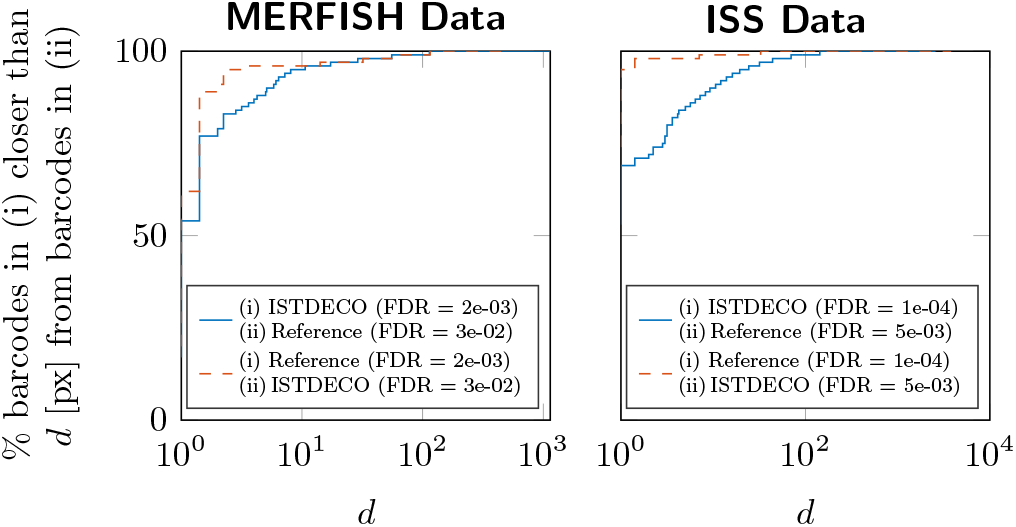
The percentage of barcodes decoded and localised by method (i) (see legend), with distance less than *d* [px] to a barcode of the same type decoded and localised by method (ii). Barcodes detected by (i) are filtered such that a low FDR is obtained, whereas barcodes detected by (ii) are filtered such that a slightly higher FDR is obtained.

## 5 Discussion & Conclusion

Here we have proposed ISTDECO — a deconvolution tool for localising and decoding barcodes in in situ spatial transcriptomic images. The tool builds on maximising the likelihood of a Poisson statistics using multiplicative updates. The update in Eq. 4 solely consists of vectorised operations and can therefore efficiently be computed with a GPU. For that matter, we chose to implement ISTDECO using the deep learning framework PyTorch. With an 8GB laptop GPU we managed to process a 2048 × 2048 field of view with 8 rounds, 2 channels and 140 barcodes in less than 30 seconds. Even larger speedup can be obtained by adopting down-sampling strategies similar to Chen *et al.* (2020).

The algorithm is relatively simple and can be implemented in just a few lines of code. Despite its simplicity, the method performs well on synthetic data, also at high signal densities, outperforming heuristic algorithms, and comparable to similar algorithms such as BarDensr.

It should also be noted that ISTDECO requires the spots to be adequately aligned across the rounds and channels. This can be achieved by registering the image data. We found a point-cloud based registration like Qian *et al.* (2020) to be very effective.

In terms of the experiment on MERFISH image data, we found a strong correlation in how frequent the different types of barcodes were detected. We also found that most of the ISTDECO detected barcodes are in proximity of similar barcodes detected by the reference algorithm. This indicates an agreement between the MERFISH reference method and ISTDECO. We also noted a strong correlation between ISTDECO and the ISS reference method. Occasionally, on the ISS images, fluorescence was not ideally washed away be tween sequencing rounds, resulting in situations where fitting a barcode became very ambiguous — even for hand-curated methods.

In conclusion, we believe that ISTDECO is a simple yet effective tool for molecular detection in multiple xed images.

## Acknowledgements

We would like to express our gratitude for the Nilsson lab for providing image data.

## Funding

This work was supported by the European Research Council [ERC-2015-CoG 682810 to C.W.]; and the Klaus Tschira foundation [Informatics for Life to F.A.H.].

## Supplementary Information

### Hyperparameter Tuning

For the speed test in Section 3.4, the parameters of ISTDECO and BarDensr were tuned using grid search. For ISTDECO this involved tuning the standard deviation *σ*_psf_ as well as the background *b*. For BarDensr we chose to only fit the rolony matrix (using the dense learner) as this would decrease the computational time during testing. The other parameters tuned were the blur level of the heat kernel, the baseline *b*, and the parameter controlling the regularisation. For both methods, the signal spots were located through non-maxima suppression using a radius of 1 [px]. The *F*_1_ score was computed after different number of optimisation iterations by searching for the intensity threshold that gave the highest *F*_1_. This search was not included in the timing. This text is supplementary to Section 3.4 in the main text.

### Supplementary Figures

**Fig. 7.**
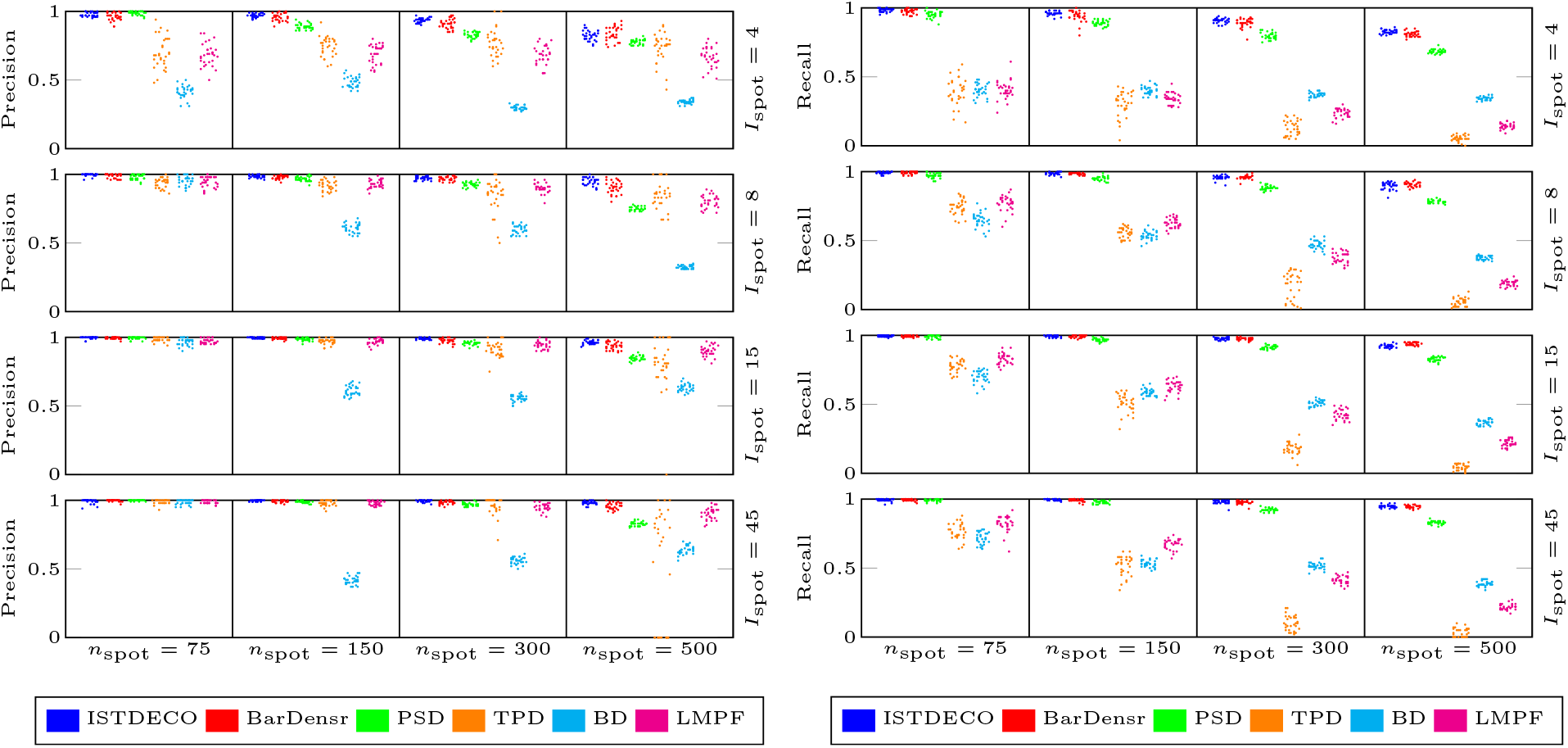
Precision (left) and Recall (right) for respective methods. Supplementary to Figure 3.

**Fig. 8.**
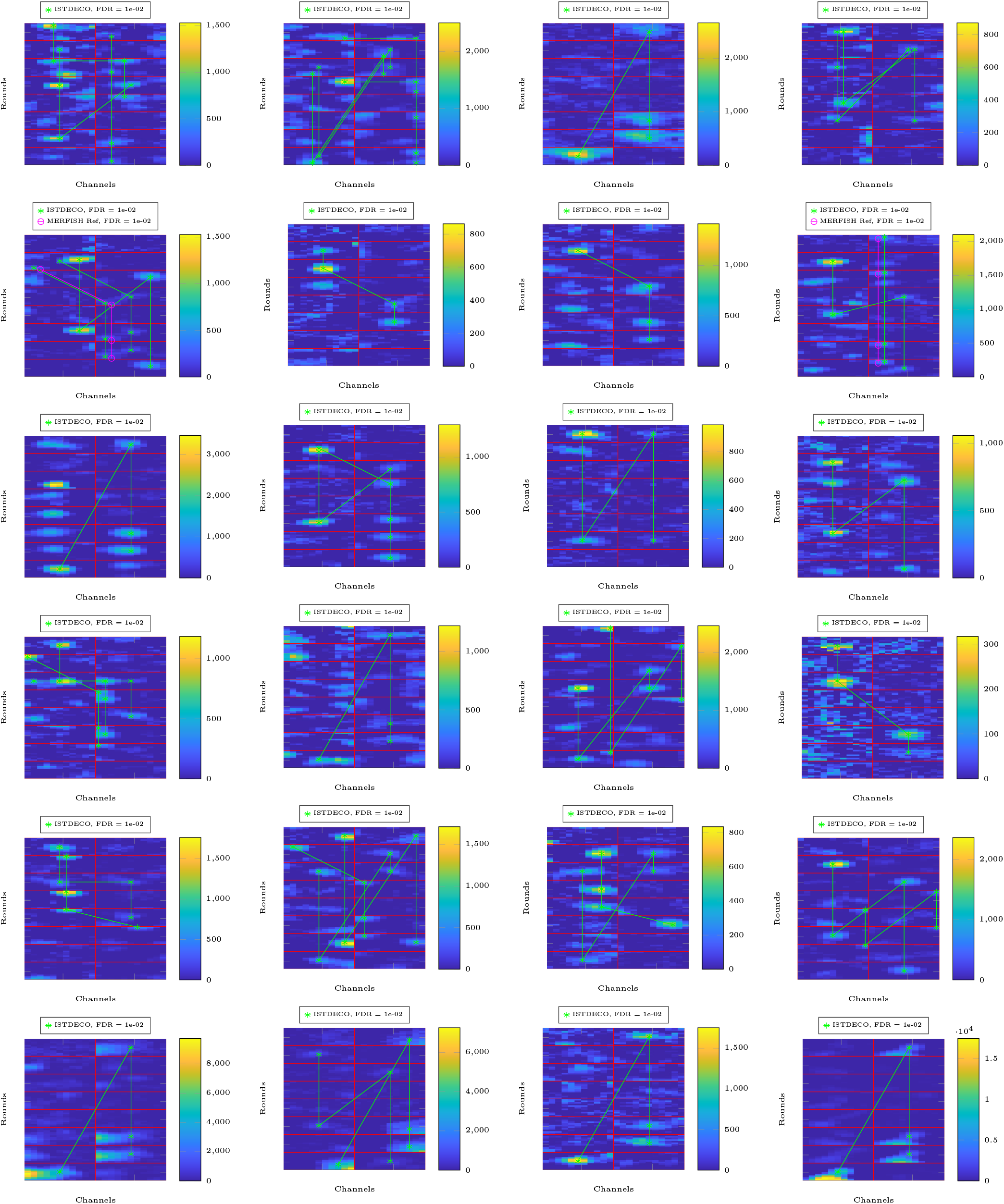
Comparison between barcodes detected by ISTDECO and barcodes in the MERFISH reference data. Detections are thresholded such that an FDR of 0.01 is obtained. Each image is centred around a barcode that is detected by ISTDECO but is not found within a radius of 3 [px] in the reference data.

**Fig. 9.**
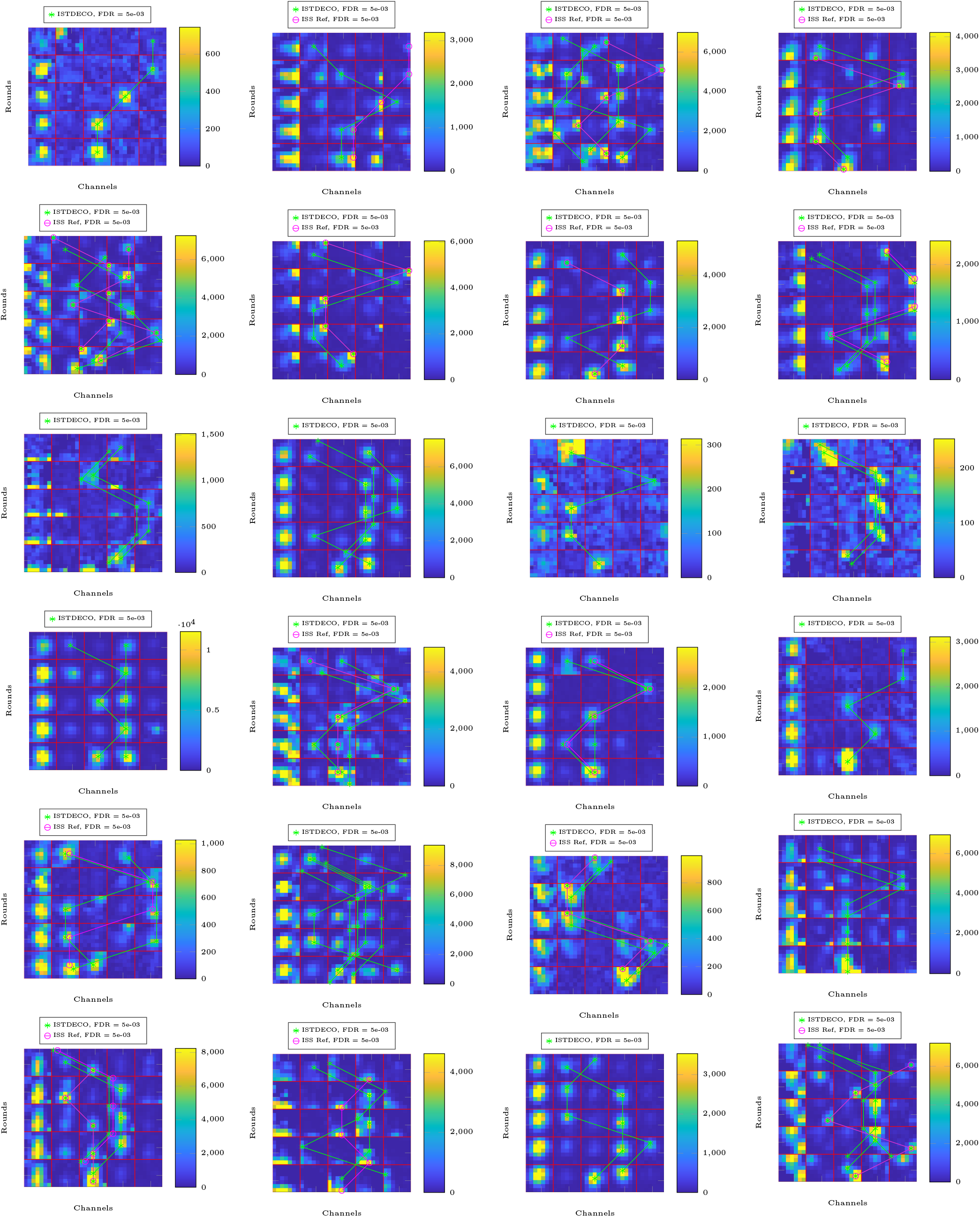
Comparison between barcodes detected by ISTDECO and barcodes in the ISS reference data. Detections are thresholded such that an FDR of 5e-3 is obtained. Each image is centred around a barcode that is detected by ISTDECO but is not found within a radius of 3 [px] in the reference data. The first channel is an *anchor* channel showing all bases.

## Notes

### Competing Interest Statement

The authors have declared no competing interest.

